## Supplementary Figures for "A therapeutic combination of two small molecule toxin inhibitors provides pancontinental preclinical efficacy against viper snakebite"

**Supplementary Information**

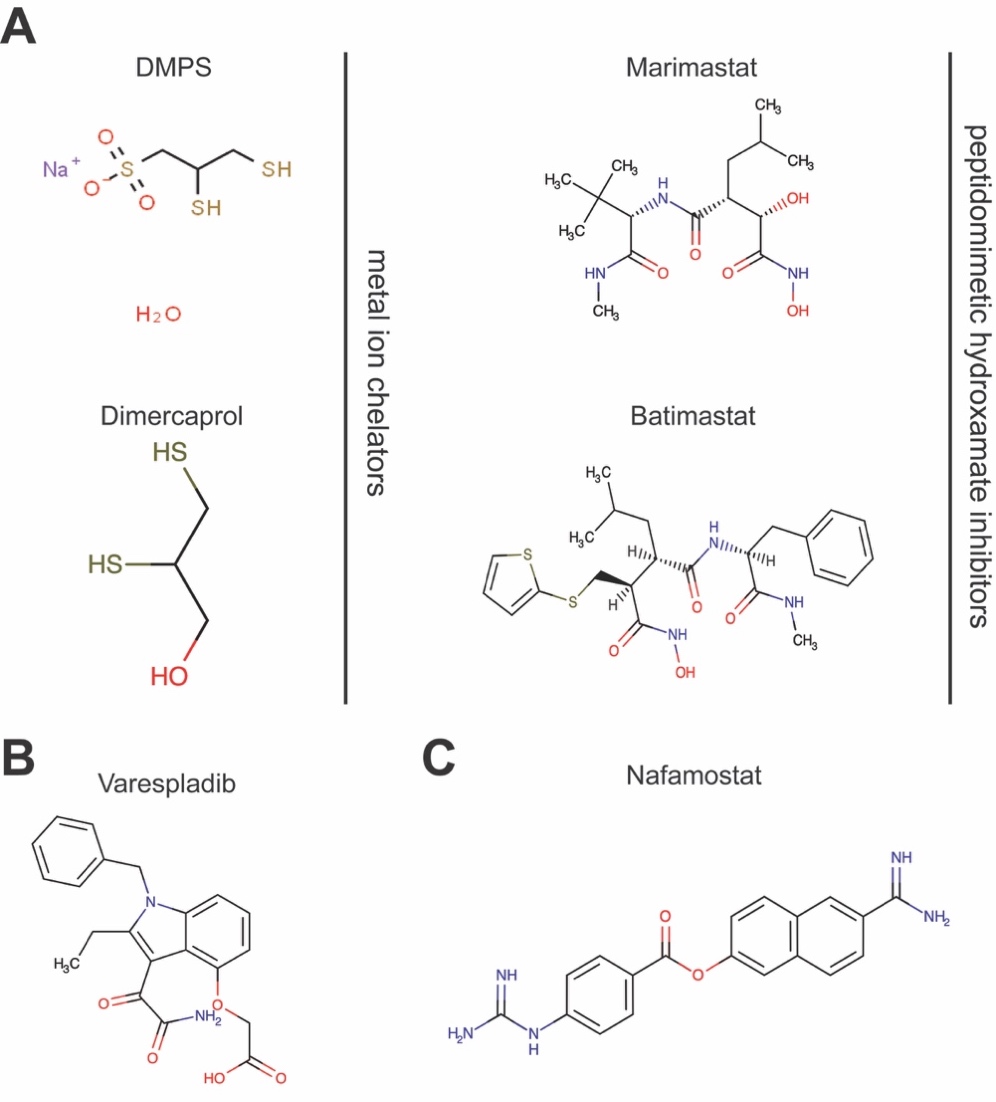

**Fig. S1. Chemical structures of the small molecule toxin inhibitors used in this study.** **(A)** Snake venom metalloproteinase (SVMP)-inhibitors: the metal ion chelators, DMPS (unithiol) and dimercaprol, and the peptidomimetic hydroxamate inhibitors, marimastat and batimastat. **(B)** The secretory phospholipase A_2_ (PLA_2_)-inhibitor, varespladib. **(C)** The serine protease-inhibitor, nafamostat.

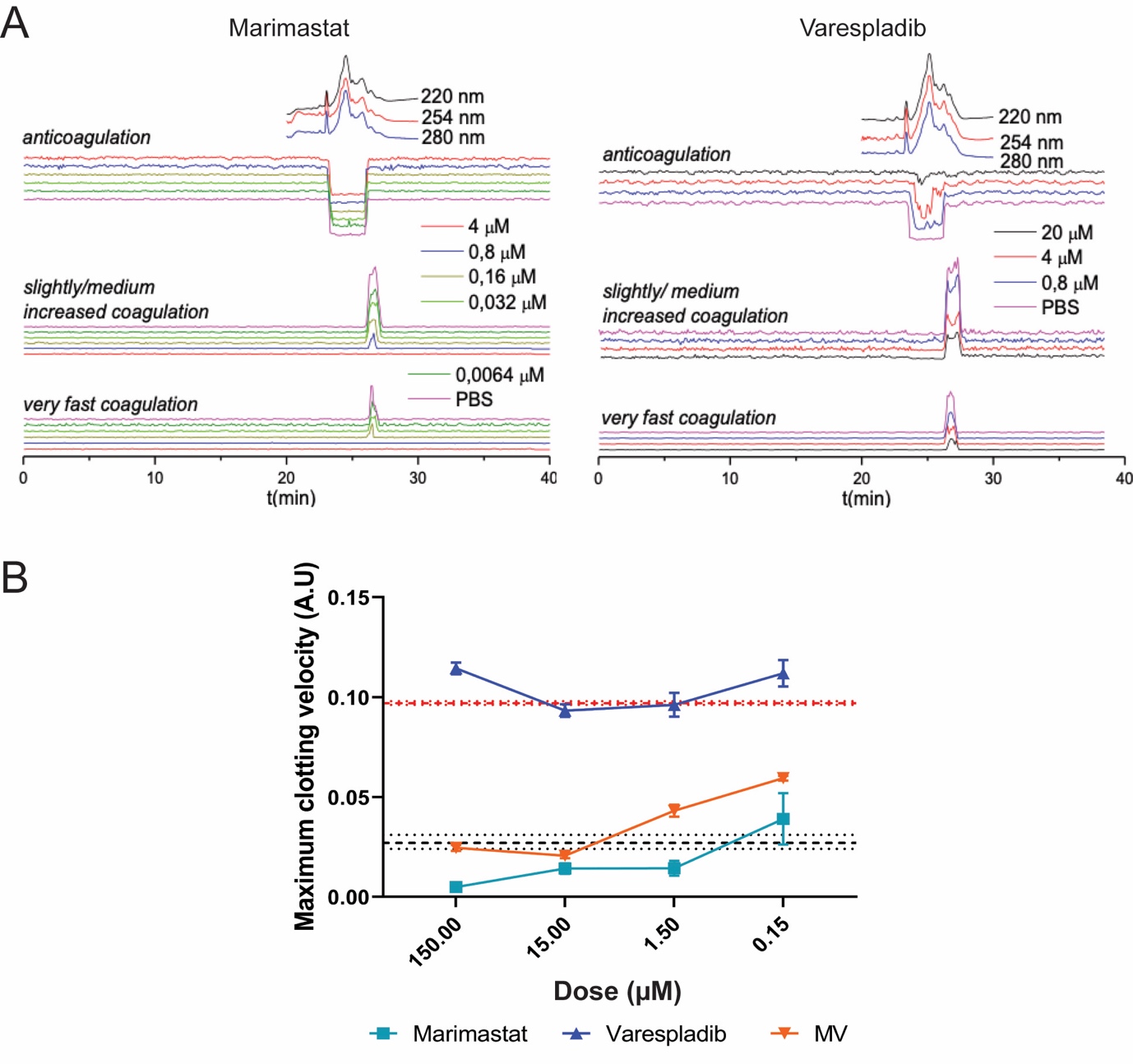

**Fig. S2. The inhibitory effects of marimastat and varespladib against procoagulant and anticoagulant toxins fractionated from Russell’s viper (*D. russelii*) venom.** **(A)** Representative nanofractionation chromatograms showing the neutralizing potency of marimastat and varespladib against the pro- and anti-coagulant activities of *D. russelli* venom, respectively. The data is plotted separately for ‘very fast coagulation’, ‘medium coagulation’ and ‘anticoagulation’, based on the slopes of the 0-5 min and 0-20 min readings, and the single endpoint reading at 180 min. The tested inhibitor concentrations are color-coded and presented alongside the chromatograms. **(B)** The interplay between marimastat and varespladib in neutralizing coagulation-related activities of *D. russelii* venom across a 150 µM-150 nM concentration range. Inhibitors are color-coded: marimastat, turquoise; varespladib, blue, marimastat + varespladib, orange. The data is presented as the maximal clotting velocity at each concentration, and represents means of triplicate independent repeats with SEMs, where each technical repeat represents the mean of n≥2 technical replicates. The venom-only control (dashed red lines) and the negative control (dashed black lines) are presented as intervals and represent the average ± SEM observed for the venom-only and PBS controls across all datasets and replicates.

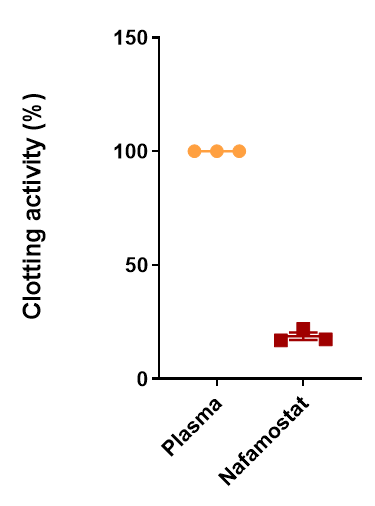

**Fig. S3. The serine protease-inhibitor nafamostat inhibits normal plasma clotting.** Comparison of normal clotting (i.e. in the absence of venom and inhibitors via recalcification) and clotting in the presence of the generic serine protease-inhibitor nafamostat. At high doses (150 µM), nafamostat dramatically inhibits plasma coagulation. The anticoagulant effect of this drug is expressed as a percentage of the plasma-only activity.

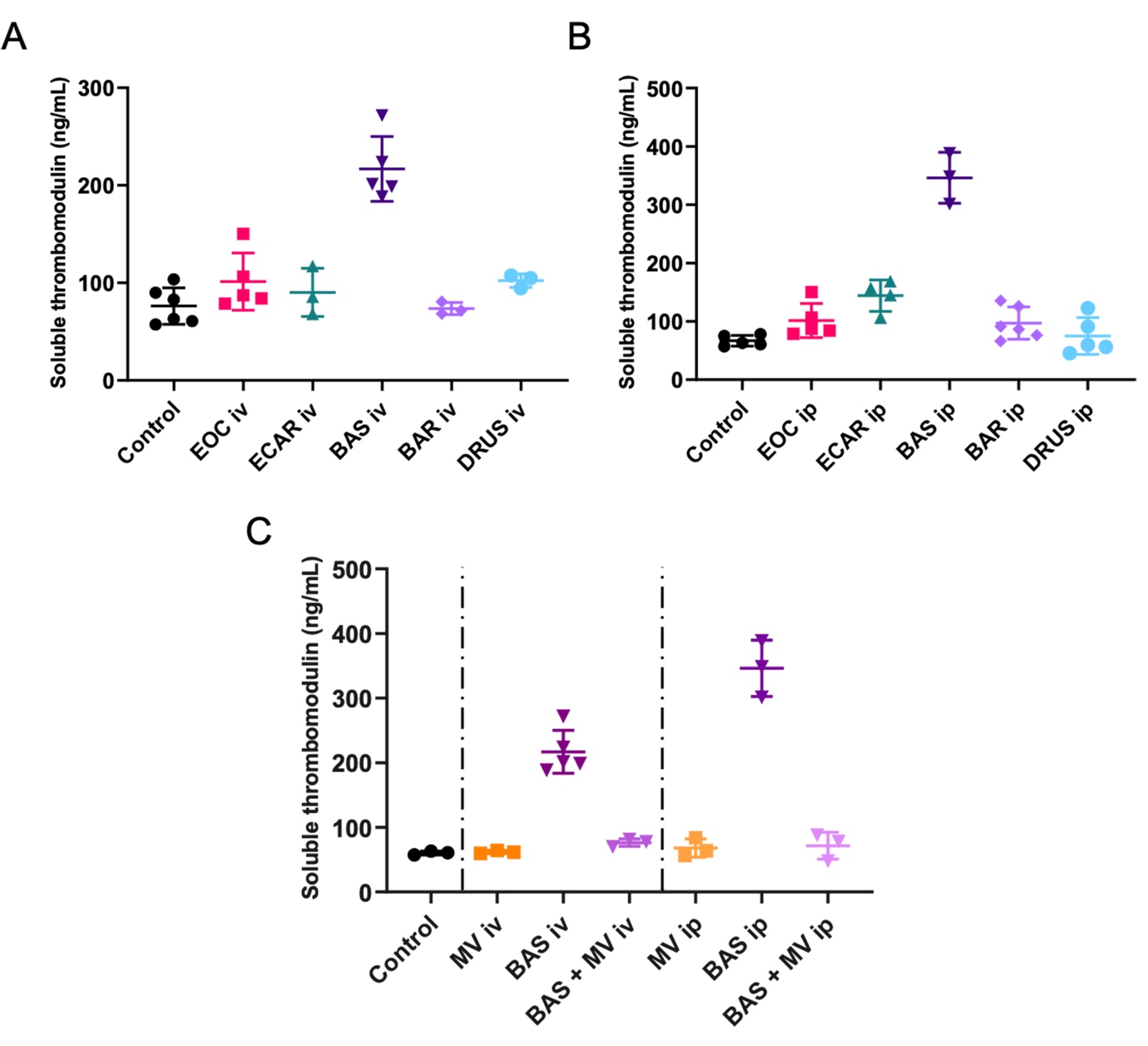

**Fig. S4. Soluble thrombomodulin is a valuable biomarker for *Bothrops asper* envenoming.** Quantified levels of soluble thrombomodulin determined by ELISA experiments using mouse plasma collected from experimental animals challenged with venoms ± inhibitors via the intravenous (iv) and intraperitoneal (ip) routes. Quantified soluble thrombomodulin levels in animals that received venom iv **(A)** and ip **(B)** reveal elevations caused by *B. asper* venom. **(C)** Treatment with the marimastat and varespladib dual combination therapy (MV) reduces *B. asper-*elevated soluble thrombomodulin levels to values comparable with the control. Where the time of death was the same within experimental groups (e.g. early deaths or complete survival) thrombomodulin levels were quantified for n=3, and where times of death varied, n=5. The data displayed represents means of the duplicate technical repeats plus SDs.

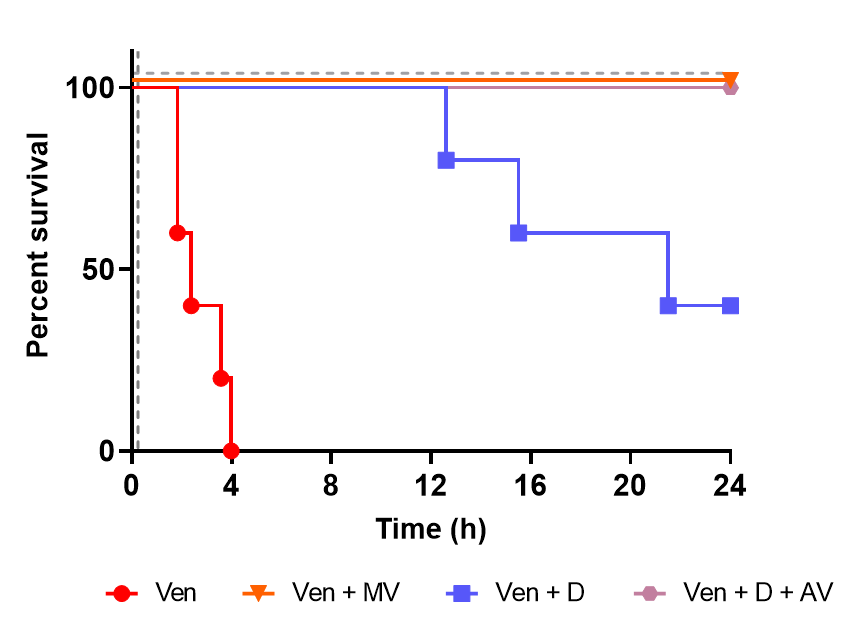

**Fig. S5. The therapeutic combination of marimastat and varespladib outperforms the licensed metal chelator DMPS, and shows equipotency with a combination of DMPS and conventional antivenom in a ‘challenge then treat’ model of envenoming.** Kaplan-Meier survival graphs for experimental animals (n=5) that received *E. ocellatus* venom (intraperitoneal administration of 90 µg; 5 × iv. LD_50_), followed by delayed drug treatment intraperitoneally 15 mins later. Drug doses were kept constant at 120 µg for each inhibitor used in the solo DMPS (D) treatment group and the marimastat and varespladib combination therapy (MV) group. For the DMPS and antivenom (D + AV) group, experimental animals received 120 µg of DMPS 15 mins after venom, and intravenous antivenom (168 µl of the *E. ocellatus* monospecific antivenom EchiTAbG, MicroPharm Ltd, UK) 1 hr after venom delivery. All DMPS data presented is taken from Albulescu et al.^28^.
